# DenoiseST: A dual-channel unsupervised deep learning-based denoising method to identify spatial domains and functionally variable genes in spatial transcriptomics

**DOI:** 10.1101/2024.03.04.583438

**Authors:** Yaxuan Cui, Ruheng Wang, Xin Zeng, Yang Cui, Zheyong Zhu, Kenta Nakai, Xiucai Ye, Tetsuya Sakurai, Leyi Wei

**Author notes:** To whom correspondence should be addressed: Xiucai Ye, Leyi Wei. Co-first author (Yaxuan Cui, Ruheng Wang).

## Abstract

Spatial transcriptomics provides a unique opportunity for understanding cellular organization and function in a spatial context. However, spatial transcriptome exists the problem of dropout noise, exposing a major challenge for accurate downstream data analysis. Here, we proposed DenoiseST, a dual-channel unsupervised adaptive deep learning-based denoising method for data imputing, clustering, and identifying functionally variable genes in spatial transcriptomics. To leverage spatial information and gene expression profiles, we proposed a dual-channel joint learning strategy with graph convolutional networks to sufficiently explore both linear and nonlinear representation embeddings in an unsupervised manner, enhancing the discriminative information learning ability from the global perspectives of data distributions. In particular, DenoiseST enables the adaptively fitting of different gene distributions to the clustered domains and employs tissue-level spatial information to accurately identify functionally variable genes with different spatial resolutions, revealing their enrichment in corresponding gene pathways. Extensive validations on a total of 18 real spatial transcriptome datasets show that DenoiseST obtains excellent performance and results on brain tissue datasets indicate it outperforms the state-of-the-art methods when handling artificial dropout noise with a remarkable margin of ∼15%, demonstrating its effectiveness and robustness. Case study results demonstrate that when applied to identify biological structural regions on human breast cancer spatial transcriptomic datasets, DenoiseST successfully detected biologically significant immune-related structural regions, which are subsequently validated through Gene Ontology (GO), cell-cell communication, and survival analysis. In conclusion, we expect that DenoiseST is a novel and efficient method for spatial transcriptome analysis, offering unique insights into spatial organization and function.

## Introduction

The spatial distribution of transcripts is crucial for understanding cellular states and cellular organization in tissues, and numerous methodologies for spatially profiling gene expression have emerged in recent years^1^. Within the realm of genomics, progress in massively parallel deoxyribonucleic acid (DNA) sequencing, molecular biology, DNA-based molecular barcoding, and computational analysis has empowered the quantification of gene expression^2^. Furthermore, these advancements have extended to the recent capability of scrutinizing epigenetic regulation in numerous individual cells^2^. These strategies and concepts were creatively adapted to locally capture single-cell ribonucleic acid (RNA) from intact tissue sections on pixelated DNA-barcoded surfaces, and to read out their genetic identities using next-generation sequencing^3^. We call this technology family “sequencing-based spatial transcriptomics” (sST)^2^, such as 10x Visium^4^, Slide-seq^5,6^, Stereo-seq^7^, and PIXEL-seq^8^ have enabled the genome-wide profiling of gene expression at captured locations (referred to as “spots”). The technological revolution in spatial transcriptomics (ST) has overcome many key limitations of single-cell ribonucleic acid sequencing (scRNA-seq)^9^. Linking cellular gene expression with its spatial distribution holds vital significance for gaining an in-depth understanding of biological functions, describing interactive biological networks, and mechanisms underlying disease development^10^. Spatial information proves valuable not only in deciphering cell-cell communications but also in unraveling gene regulatory networks^11^. The production of extensive volumes of spatial transcriptome data dictates that computational efforts go hand-in-hand with experimental methods^12^. A fundamental component of spatial transcriptome profiling analysis involves clustering spots to unveil cell types and deduce cell lineages through the examination of transcriptome relationships among cells^13,14^. Unsupervised clustering is crucial for the analysis of spatial transcriptome data to identify novel cell types. Existing algorithms that identify clusters in spatial domains, such as k-means, Louvain, and Seurat, use only gene expression data to cluster cells at different locations (spots) into their respective spatial domains^15^. The spatial domains typically identified by these methods exhibit discontinuity, as they tend to underutilize spatial information in discerning spot locations that potentially belong to the same spatial domain^16,17^.

Recent spatial transcriptome clustering methods^18^ take into account the similarity between adjacent spots, aiming to better capture the spatial dependence of gene expression. For example, BayesSpace is a Bayesian statistical method that encourages neighboring spots to belong to the same cluster by introducing spatial neighbor structure into the prior^19^. SpaGCN applies a graph convolutional network to integrate gene expression and spatial location, with the additional integration of a self-supervised module for identifying spatial domains^20^. STAGATE utilizes a graph-attention autoencoder framework to identify spatial domains by integrating spatial information and gene expression profiles^21^. DeepST uses a variational graph autoencoder framework that integrates spatial location, histology, and gene expression to model spatially embedded representations for identifying spatial domains with similar expression patterns and histology^22^. GraphST is a method for graph self-supervised contrastive learning, which fully utilizes spatial information and gene expression profiles for spatial information clustering^23^.

Previous studies have compared and summarized the performance of multiple tools, and nearly no clustering method has robust performance and clusters well for all spatial transcriptome datasets^18^. In addition, another challenge in whole-genome spatial transcriptome data lies in the presence of technical noise during sequencing and the abundance of zeros in the spatial transcriptome data^24^ (Zeros often make up more than 50% of the total genes with expression, commonly referred to as dropout noise.). Dropout noise is frequently induced by low RNA capture rates, leading to the occurrence of false zero counts in gene expression levels^25^. Researchers have different views on the larger number of high zero values in data based on next-generation sequencing^26^. Some believe that zero represents no expression or low expression of biological signals^27,28^, while others believe that zero indicates missing data that needs correction^29–32^. A recent study shows that the generation mechanism of non-biological zeros is protocol dependent, and the Poisson, zero-inflated Poisson (ZIP), and negative binomial (NB) models are special cases of the zero-inflated negative binomial (ZINB) model^33^. The deep learning model of ZINB demonstrates superior performance in single-cell imputation^30^. Furthermore, determining the optimal number of clusters in the absence of data label is also a challenge in the clustering process of spatial transcriptome data, as it significantly affects the accuracy and practicality of the clustering^34,35^. All of these are aspects that need careful consideration.

Identifying marker genes using spatial information is another crucial problem in spatial transcriptomics^36,37^. Recognizing genes with rich expression in the spatial domain is of paramount importance. Currently, numerous tools have been developed for detecting spatially variable genes, such as Trendsceek^38^, SpatialDE^39^, SPARK^40^, SpaGCN^20^, and nnSVG^41^. However, firstly, some methods overlook the differential distribution of different genes. Secondly, some of them might neglect the prior knowledge at the cluster level, while others may excessively rely on cluster-level priors, thereby disregarding differences between clustering results and the true ground truth. These challenges contribute to the potential issue of false positives in the identification of spatially variable genes. Furthermore, there is an urgent need to address the identification of genes that not only conform to spatial distribution but also exhibit pathway enrichment related to relevant tissues. This is crucial to assist researchers in gaining a more in-depth understanding of specific tissues, with accurate annotation of the ground truth.

Based on these observations, we developed DenoiseST, a dual-channel unsupervised deep learning-based denoising method that leverages spatial information and gene expression profiles to overcome the impact of dropout noise for spatial transcriptome clustering and identification of functionally variable genes. In particular, we proposed an adaptive joint learning strategy with consensus clustering techniques and graph convolutional networks to obtain linear and nonlinear representations respectively in an unsupervised manner. Additionally, DenoiseST can automatically estimate the number of clustering clusters and adaptively construct a modeling pipeline based on user-provided data. Adaptive modeling is beneficial for accommodating diverse data and exhibiting generalization, thereby preventing model overfitting or underfitting. To address the limitations in the Identification of spatially variable genes, we proposed functionally variable genes. Analyzing the identified functionally variable genes, we observed their favorable spatial continuity on slices, and enrichment analysis validated they are related to certain functional pathways. Applying this method to 12 human dorsolateral prefrontal cortex (DLPFC) datasets generated from spatial transcriptome, we found that DenoiseST outperforms other state-of-the-art methods by a significant margin in terms of cluster accuracy, under the scenarios either with known cluster numbers or with estimated cluster numbers. DenoiseST clusters high-resolution spatial transcriptomic data, revealing contiguous regions with tissue hierarchy resembling that observed in tissue slices. For further evaluating our ability to accurately identify the biological structural regions, the DenoiseST was applied to two spatial transcriptome datasets from human breast cancer. The results demonstrate that DenoiseST effectively identified key biological structures, which were further confirmed by a series of analyses such as Gene Ontology (GO), cell-cell communication, and survival analysis. Finally, we discussed the robustness of the proposed DenoiseST under random parameter initialization and its ability to resist pseudo dropout noise. In summary, our results indicate that DenoiseST excels in accurately identifying spatial domains, resisting dropout noise, demonstrating model robustness, and possessing a powerful ability to accurately identify biological structures.

## Results

### Overview of the proposed DenoiseST

DenoiseST primarily performs spatial data imputing, clustering, and identifying functionally variable genes on spatial transcriptome data. It can adaptively model and conduct fully automated clustering analysis based on the input spatial transcriptome data (workflow shown in **Figure 1**). The following provides an overview of the clustering and imputation model, as well as the identifying functionally variable genes.

**Figure 1.**
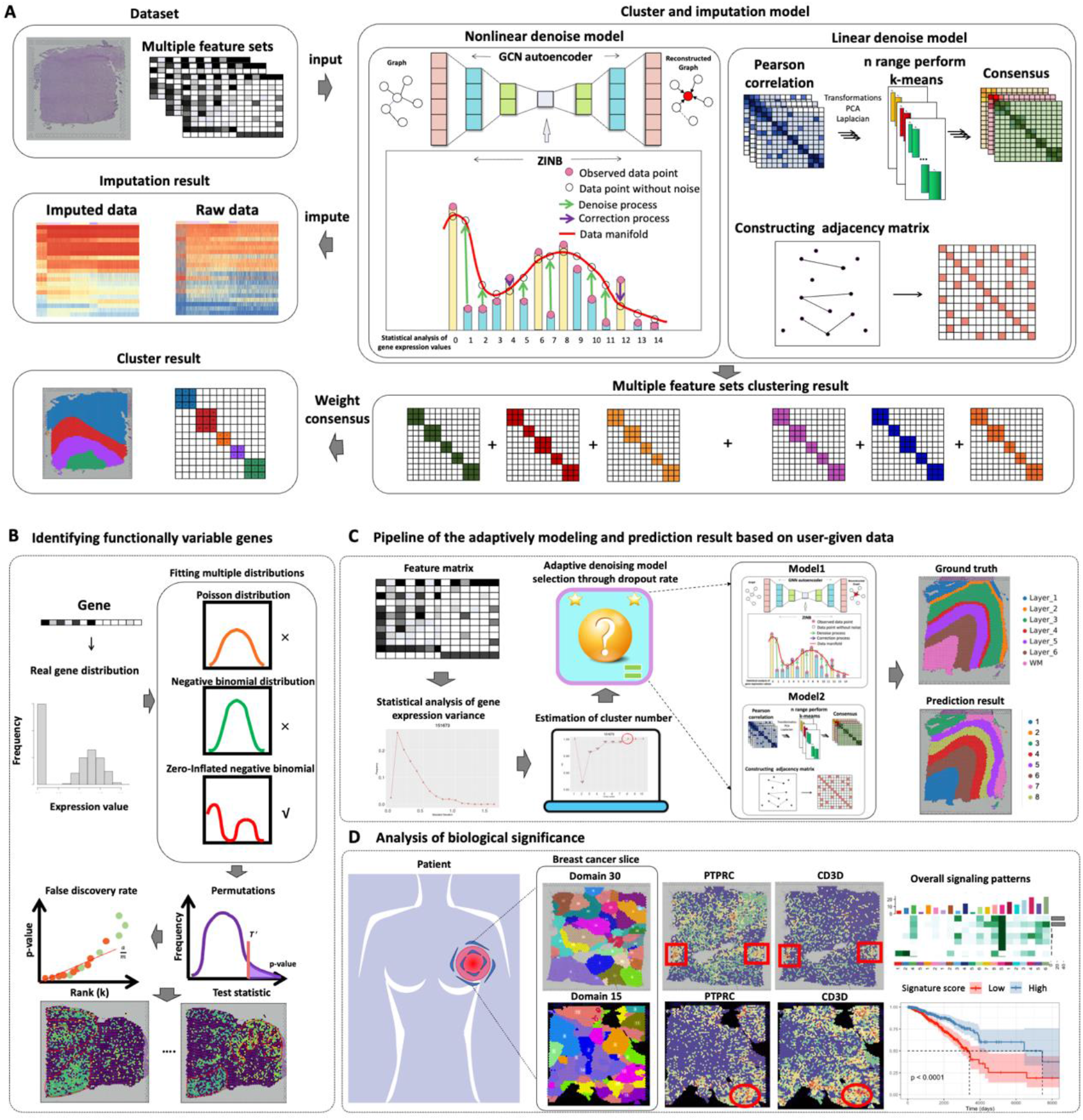
Overview of DenoiseST. **(A)** DenoiseST takes the preprocessed spatial transcriptome feature sets and a neighborhood graph constructed using spot coordinates (x, y) as input data. The data is then input into both the linear denoise model and the nonlinear denoise model that we have built. The linear denoise model utilizes transformations and incorporates spatial position information against linear dropout noise. On the other hand, the nonlinear denoise model employs a zero-inflated negative binomial (ZINB) and graph convolutional network (GCN) to address nonlinear dropout noise. Notably, the ZINB loss function of the nonlinear denoise model includes an imputation function for the original gene expression matrix. Finally, a weight consensus clustering is performed to integrate the multiple clustering results generated by both the linear denoise model and the nonlinear denoise model, respectively. **(B)** Identifying functionally variable genes. To identify functionally variable genes, we can fit the data to three commonly used distributions: the negative binomial distribution, the zero-inflated negative binomial distribution, and the Poisson distribution. Subsequently, permutation tests are conducted to calculate p-values, followed by the incorporation of histological information for selection. **(C)** Pipeline of the adaptively modeling and prediction result based on user-given data. For user-provided data, our DenoiseST firstly estimates the number of clustering clusters and the dropout rate of the data, and then adaptively chooses the appropriate model architecture to accommodate diverse data. **(D)** Analysis of biological significance. DenoiseST can be applied to human breast cancer spatial transcriptomic datasets to find biologically significant immune-related structural regions.

### Overview of clustering and imputation model

The DenoiseST model’s specific process (see **Figure 1A** and **Supplementary Figure S1**) comprises several steps. Firstly, the input data includes a gene expression matrix and spatial location information. Within the gene expression matrix, highly variable genes are identified using the ‘seurat_v3’ function. Subsequently, multiple top feature gene sets are selected and normalized. Next, the ClusterR package is employed to automatically estimate the number of clusters, based on the top 5000 features. These feature gene sets are then separately input into both the linear denoise model and the nonlinear denoise model.

In the linear denoise model, we first construct Pearson’s cell-cell similarity matrix and then perform transformations. After executing the transformations, we conduct k-means clustering on the first d eigenvectors of the transformed distance matrix to obtain the consensus matrix. This enhances the similarity of adjacent spots by iterating the adjacency matrix of spatial position information. The similarity-enhanced matrix obtained after enhancing the similarity of adjacent spots is subjected to spectral clustering methods. For the similarity-enhanced matrix, clusters are detected by a spectral clustering method.

In the nonlinear denoise model, the preprocessed spatial gene expression matrix and the spatial location information of the spatial transcriptome are used as input. The latent representation is learned using a zero-inflated negative binomial (ZINB) and graph convolutional network model^42^ (GCN), which can not only preserve informative features from gene expression profiles, spatial location information, and local contextual information but also impute the zero values of the original data to compensate for the negative impact of dropout noise in the original data. We obtain the reconstructed gene expression matrix. During our analysis of the reconstruction matrix, we discovered its function of imputation. Subsequently, we utilized the R package ‘mclust’ to cluster the reconstructed gene expression matrix.

The clustering results obtained by inputting multiple feature gene sets into the linear denoise model and the nonlinear denoise model, respectively, are merged to calculate the weight consensus clustering matrix. Finally, spectral clustering is performed on the weight consensus clustering matrix to obtain the final clustering result. Before performing clustering, the model can estimate the number of clusters for spatial transcriptomic data. Moreover, DenoiseST can adaptively choose different models for clustering based on the dropout rate in the data.

### Overview of identifying functionally variable genes

Based on the final clustering results and fitting the data to three commonly used distributions (negative binomial distribution, zero-inflated negative binomial distribution, and Poisson distribution), we calculated the best-fit distribution for the two experimental conditions. Subsequently, we computed the Bhattacharyya distance between the distributions, and to validate its significance, we performed a permutation test. We conducted permutation tests to calculate p-values, aiming to gain confidence in the Bhattacharyya distance scores. The objective of these analyses was to identify differential genes.

To identify more significant differential genes, we performed the following procedure and defined them as functionally variable genes. Firstly, we took the intersection of the differential genes defined by each cluster. Subsequently, we mapped the expression of these genes to each spot in the spatial organization and selected the 100 spots for each gene with the highest expression. Afterward, we calculated the distance between each spot and summed the number of the spots if the distance was less than a threshold. If the numerical value is higher, we believe it is more spatially continuous and thus more likely histologically related and considered a functionally variable gene (as shown in **Figure 1B** and **Supplementary Figure S2**).

### DenoiseST outperforms the state-of-the-art methods on 12 benchmark spatial transcriptomics datasets

To measure the effectiveness of DenoiseST in identifying spatial domains, we undertook a comparative analysis of DenoiseST against five state-of-the-art methods (i.e. BayesSpace, SpaGCN, DeepST, STAGATE, and GraphST) on 12 benchmark spatial transcriptomics datasets. The compared methods (Details can be seen in **Supplementary Table S2**) were executed with default parameter settings, and the performance is quantitatively measured with the ARI scores as shown in **Figure 2A** and **Supplementary Figure S3**. First, we observed that DenoiseST achieved the highest ARI score in eight datasets, while GraphST acquired the top position in four datasets, STAGATE in two datasets, and SpaGCN in one dataset. As shown in **Figure 2A**, the boxplots illustrating the ARI scores of the six methods applied to the 12 DLPFC slices reveal the following ranges: DenoiseST achieved ARI scores between 0.44 and 0.86, GraphST obtained ARI scores ranging from 0.43 to 0.63, STAGATE displayed ARI scores between 0.33 and 0.60, DeepST recorded scores ranging from 0.32 to 0.57, SpaGCN exhibited ARI scores between 0.30 and 0.57, and BayesSpace recorded scores ranging from 0.30 to 0.55. It can be concluded that DenoiseST can obtain better clustering results on 12 datasets and achieve higher ARI on 151671 (ARI=0.86) and 151672 slices (ARI=0.79). Specific clustering visualization results (**Figure 2B, Figure 2C**, and **Supplementary Figure S3**).

**Figure 2.**
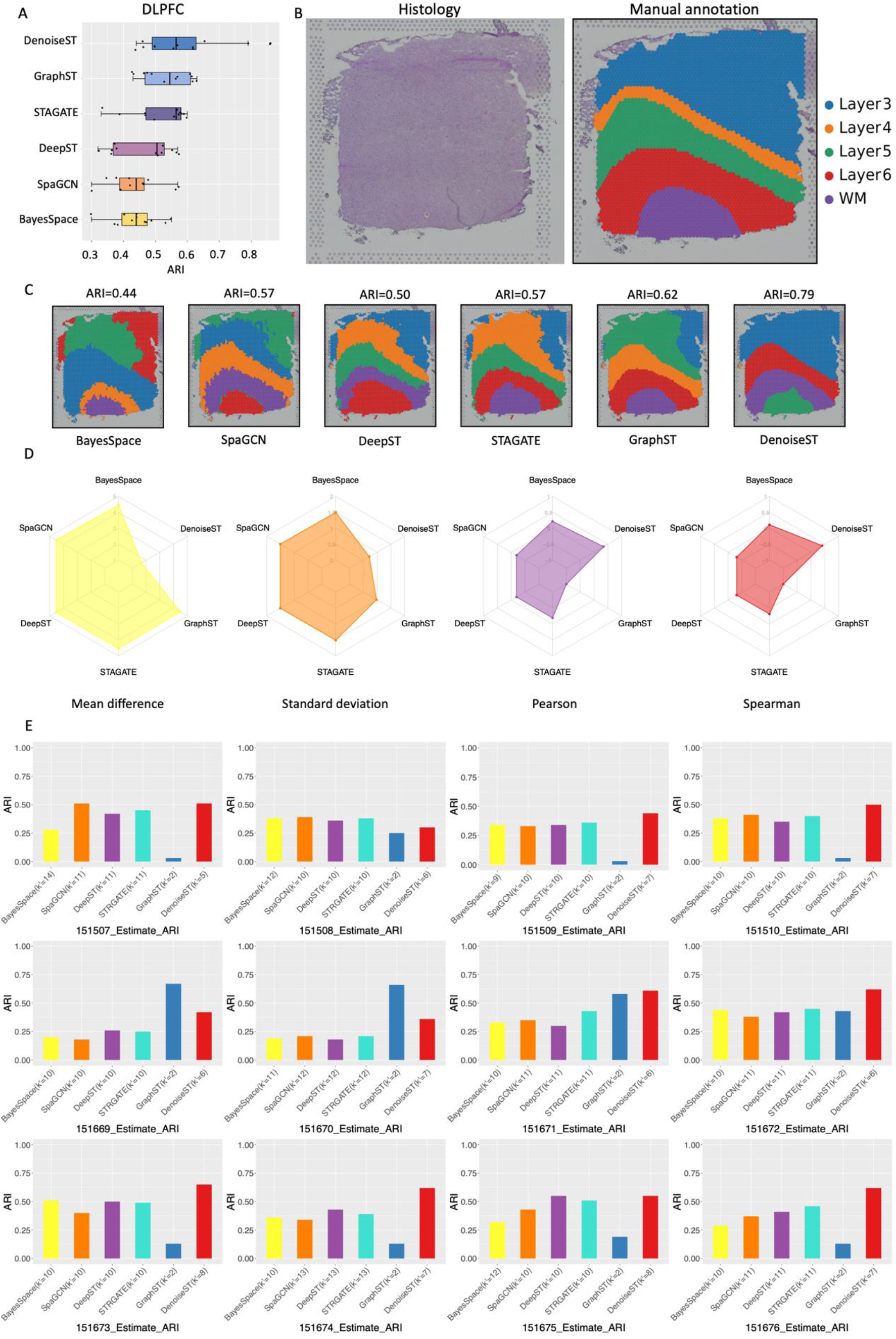
DenoiseST clustering enhances the identification of tissue structures in DLPFC. **(A)** Boxplots depicting the ARI scores of the six methods applied to the 12 DLPFC slices. The central line represents the median, the box boundaries indicate the upper and lower quartiles and the whiskers represent the 1.5 times interquartile range. **(B)** Histology and manual annotation images from the original study. **(C)** Clustering results produced by different spatial transcriptomics methods, such as BayesSpace, SpaGCN, DeepST, STAGATE, GraphST, and DenoiseST, for slice 151672 of the DLPFC dataset. Manual annotations and clustering results of the other DLPFC slices are shown in **Supplementary Figure S5. (D)** Compare clustering performance based on the estimated number of clusters. (Mean difference = 4.33, SD = 1.50, Pearson = 0.22 and Spearman = 0.11 for BayesSpace, Mean difference = 4.42, SD = 1.51, Pearson = -0.19 and Spearman = -0.31 for SpaGCN, Mean difference = 4.42, SD = 1.51, Pearson = -0.19 and Spearman = -0.31 for DeepST, Mean difference = 4.42, SD = 1.51, Pearson = -0.19 and Spearman = -0.31 for STAGATE, Mean difference = 4.33, SD = 0.98, Pearson = -1 and Spearman = -1 for GraphST, Mean difference = 0.83, SD = 0.72, Pearson = 0.35 and Spearman = 0.41 for DenoiseST). **(E)** Compare clustering performance based on the estimated number of clusters.

As an unsupervised clustering method, DenoiseST offers the flexibility to accept either the user-specified number of clusters or automatically estimate the potential number of cell clusters. Because in the real experimental process, many data sets do not provide the number of clusters. After applying DenoiseST to 12 datasets (**Supplementary Figure S4**), we found that the number of clusters estimated by DenoiseST and the number of cell types labeled by the authors were similar (P-value = 0.17, chi-squared testing), and their differences were very small (mean = 0.83 and SD = 0.72). We also utilized the other five methods (BayesSpace, SpaGCN, DeepST, STAGATE, and GraphST) to estimate the number of clusters on 12 datasets, employing their default or estimate methods as specified in the paper. We then compared these methods (**Figure 2D**). DenoiseST exhibited lower differences in mean difference and standard deviation, along with higher similarities in the Pearson Correlation Coefficient and Spearman Rank Correlation Coefficient. These results indicate that DenoiseST is more accurate in estimating the number of clusters. Subsequently, we performed clustering on the benchmark data using the estimated number of clusters obtained by the respective algorithms and compared the ARI scores. Notably, we found that DenoiseST ranked first in the ARI score for 9 datasets in **Figure 2E**. Hence, the Adjusted Rand Index (ARI) values for the clustering results obtained by DenoiseST, even when the number of slice clusters is unknown, are the highest.

This underscores the robustness of our model in dealing with scenarios where the true number of clusters is not provided.

### Integrating linear and nonlinear denoising enhances the clustering performance

How to define the similarity between spots is a technical challenge for unsupervised spatial transcriptome data clustering analysis. If we know the marker genes of different spot types in advance, we can obtain higher clustering accuracy. However, most spatial transcriptome datasets have no such marker genes available. To solve this problem, most spatial transcriptome clustering algorithms preprocess the data to select highly variable genes, and most of them use the ‘seurat_v3’ method to identify the top 3000 highly variable genes. The ‘seurat_v3’ method is used to obtain the mean-variance relationship from the data, and we analyze the distribution curve of the standard deviation (SD) on the 12-slice dataset using the ‘seurat_v3’ method (as illustrated in **Supplementary Figure S5**). For a given dataset, we found that the top 5000 highly variable genes (genes with a significant level, most of which have a variance value greater than 0) can be used as a candidate gene pool. Through our analysis of spatial transcriptome data based on next-generation sequencing methods (as shown in **Supplementary Table S1** and **Supplementary Note S1**), another important factor to consider is reducing the negative impact of dropout noise.

We input multiple feature sets into the linear denoise model and the nonlinear denoise model, respectively. To eliminate the influence of linear noise, we enhance the similarity of adjacent spots in the consensus matrix. In the linear denoise model, mitigating the adverse effects of linear noise is achieved by processing the transformed metric matrix. Subsequently, we enhance the similarity of adjacent spots in the metric matrix. Moreover, we integrate adjacent point features by employing a combination of ZINB loss and a GCN model, aiming to address the impact of nonlinear noise. We analyzed the clustering results of the linear denoise model and the nonlinear denoise model on 12 spatial transcriptome datasets. We calculated the Adjusted Rand Index (ARI) score, which is widely used for cluster analysis in spatial transcriptome data. We analyzed the clustering outcomes achieved by the linear denoise model and the nonlinear denoise model on two spatial transcriptome datasets, employing the Adjusted Rand Index (ARI) score for evaluation. The ARI score is a widely adopted metric for assessing cluster analysis performance in the context of spatial transcriptome data.

We conducted testing on the linear denoise model using two low-resolution slice datasets, denoted as 151672 and 151673. The evaluation involved assessing the variations in ARI score across different feature numbers and different radius sizes for the number of spot neighbors. Next, we tested different feature numbers and different numbers of nearest spot neighbors on the nonlinear denoise model to assess its ARI score (as shown in **Figure 3B** and **Figure 3F**). We analyzed a total of 12 datasets (refer to **Supplementary Figure S6** and **Supplementary Figure S7**). Our findings indicate that the linear denoise model exhibited enhanced stability in results when employing the top 4000, 4500, and 5000 features with a radius set to 4 in these datasets. As well as the nonlinear denoise model consistently produced stable results when utilizing the top 4000, 4500, and 5000 features, with the number of neighbors set to 4.

**Figure 3.**
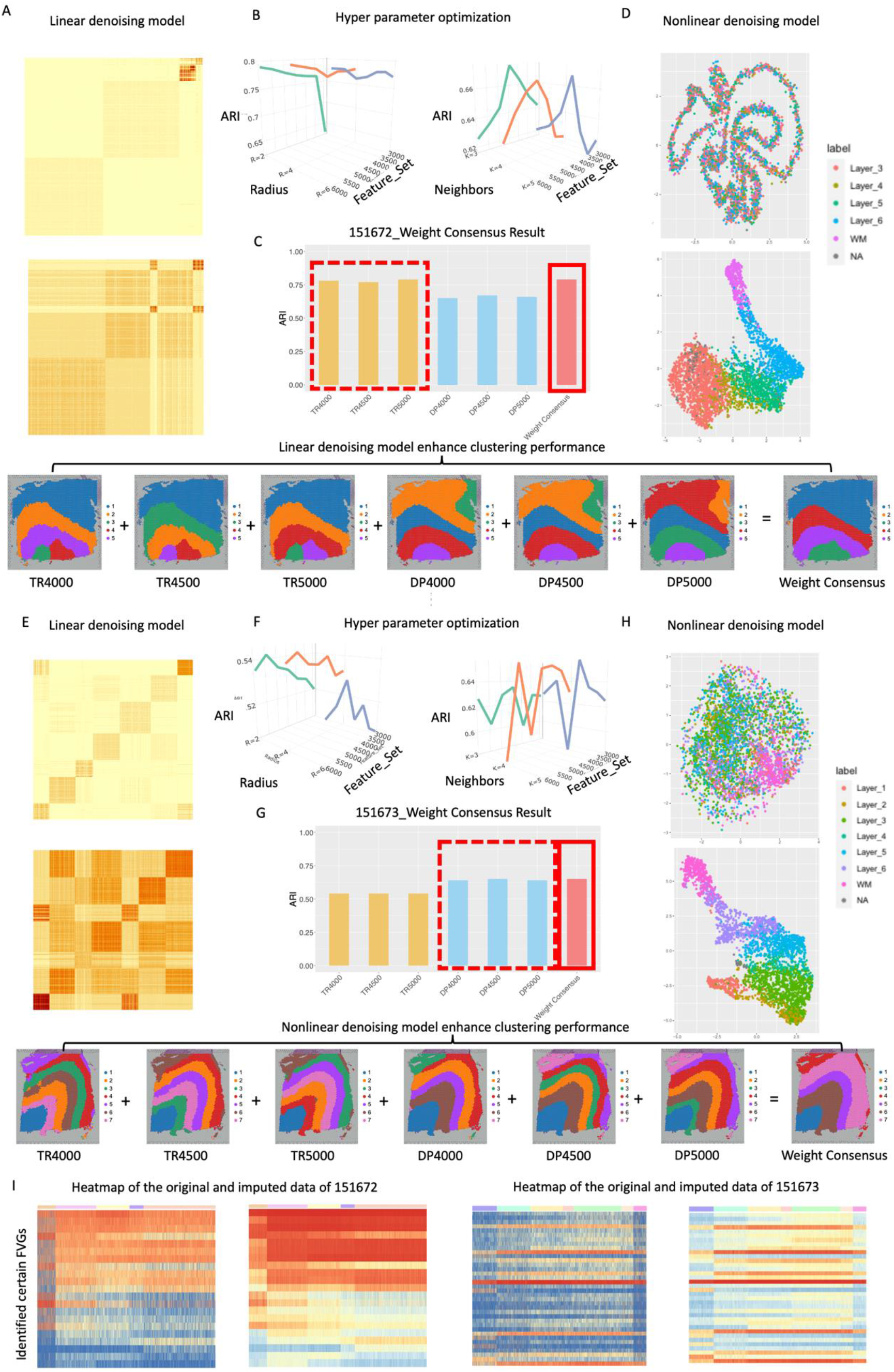
Analytical investigations are conducted within the DenoiseST model on slices 151672 and 151673, respectively. **(A)** Visualization of the original metric matrix and the consensus metric matrix of the top 5000 features in the linear denoise model of 151672. **(B)** ARI scores in the linear denoise model and the nonlinear denoise model are obtained by varying the number of features and neighbors in the adjacency matrix of 151672. **(C)** the ARI scores were calculated for six feature sets individually, and the weight consensus clustering results were based on the combination of six feature sets of 151672 (TR represents the resulting output by the linear denoise model, and DP represents the resulting output by the nonlinear denoise model). **(D)** A two-dimensional scatter plot of the original top 5000 features and reconstructed top 5000 features in the nonlinear denoise model of 151672. **(E)** Visualization of the original metric matrix and the consensus metric matrix of the top 5000 features in the linear denoise model of 151673. **(F)** ARI scores in the linear denoise model and the nonlinear denoise model are obtained by varying the number of features and neighbors in the adjacency matrix of 151673. **(G)** the ARI scores were calculated for six feature sets individually, and the weight consensus clustering results were based on the combination of six feature sets of 151673 (TR represents the resulting output by the linear denoise model, and DP represents the resulting output by the nonlinear denoise model). **(H)** A two-dimensional scatter plot of the original top 5000 features and reconstructed top 5000 features in the nonlinear denoise model of 151673. **(I)** Shows heatmaps of the original gene expression data and the reconstructed gene expression data of 151672 and 151673.

To enhance the optimization of defining feature sets, we employed multiple feature sets for weight consensus clustering, as detailed in the Materials and Methods section. In the two slice datasets, 151672 and 151673 (refer to **Figure 3C** and **Figure 3G**), we observed that the linear denoise model, employing traditional machine learning, exhibited superior clustering results on the 151672 slice. Conversely, on the 151673 slice, the nonlinear denoise model based on deep learning demonstrated better clustering outcomes. Moreover, the adverse effects of negative clustering results were mitigated through the application of weight consensus clustering.

We further analyzed 12 benchmark low-resolution slice data sets, we found that the ARI scores from weight consensus clustering were generally higher than those using a single feature set (as shown in **Supplementary Figure S8**). The weight consensus clustering results of 8 slice datasets (151507, 151510, 151670, 151671, 151672, 151673, 151675, 151676) achieved the highest ARI score. While the weight consensus clustering results for the other datasets (151508, 151509, 151669, and 151674) are not maximum with DenoiseST, the ARI score can remain at a high level. To further underscore the importance of the linear denoise model in improving the similarity between adjacent spots in the metric matrix and the nonlinear denoise model in integrating adjacent spot features. In **Figure 3A** and **Figure 3E**, we use the 151672 and 151673 slices to display the consensus metric matrix constructed using the top 5000 features of the linear denoise model, along with the metric matrix that enhances the similarity of adjacent spots. It can be observed that enhancing the similarity of adjacent spots in the consensus matrix exhibits darker colors within each cluster, and the separation between different clusters becomes increasingly clearer. Therefore, it can be proven that fusing the adjacency matrices of adjacent spots further enhances the relative affinity between spots of the same cluster (All 12 benchmark low-resolution slices are visually presented in the linear denoise model within the **Supplementary Figure S9**). In **Figure 3D** and **Figure 3H**, the nonlinear denoise model demonstrates that by fusing the top 5000 features with the adjacent spot features, UMAP dimensionality reduction is performed to create a two-dimensional plot. Compared to the original matrix, this plot more clearly shows tightly clusters, thereby enhancing the similarity of features within the same cluster (Visual representations of all 12 benchmark low-resolution slices in the nonlinear denoising model are available in **Supplementary Figure S10**). **Figure 3I** presents a heatmap comparing the reconstructed matrix output by the nonlinear denoise model with the original data matrix for the identified marker genes. We observed a significant amount of dropout noise in the original data matrix, which blurs the cell type identity. However, the reconstruction matrix more accurately portrays the differential expression of specific gene subsets, facilitating the identification of distinctions between tissues. This underscores our model’s capacity to impute the original data.

### DenoiseST has robustness against random initialization and dropout noise

Many models and tools based on machine learning and deep learning exhibit high sensitivity to both data and model parameter initialization. Varied initialization can lead to divergent outcomes. To assess the stability of the model more comprehensively, we performed 10 random clustering analyses on 12 datasets using six methods (DenoiseST, GraphST, STAGATE, DeepST, SpaGCN, and BayesSpace), as shown in **Figure 4A** and **Supplementary Figure S11**. We found that DenoiseST ranks first in median ARI score on 8 datasets, GraphST ranks first in median ARI score on 1 dataset and STAGATE ranks first in median ARI score on 3 datasets. Furthermore, our investigation unveiled that DenoiseST demonstrated high stability in 8 dataset slices, while GraphST exhibited high stability in 5 dataset slices. STAGATE displayed high stability in 2 dataset slices. Additionally, SpaGCN and BayesSpace both demonstrated high stability on 2 dataset slices each (SD<0.3).

**Figure 4.**
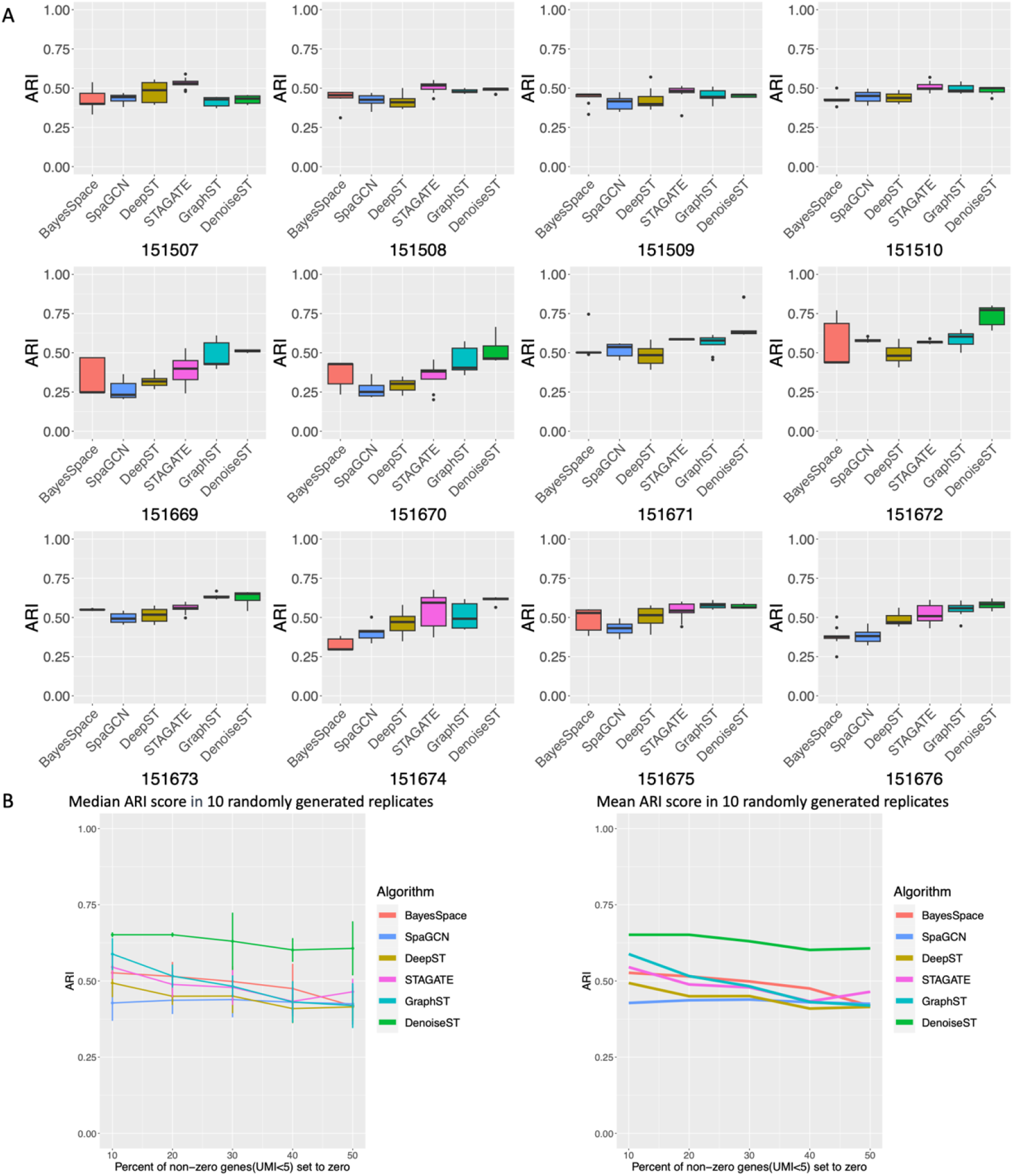
Stability comparison of DenoiseST and other methods and robustness against dropout noise. **(A)** The ARI variations of random initialization of DenoiseST, GraphST, STAGATE, DeepST, SpaGCN, and BayesSpace. **(B)** Performance comparisons of six methods on different percentages of artificial dropout noise added to the 151672 slice dataset.

To test the effectiveness of DenoiseST in mitigating the impact of dropouts, we conducted tests on the 151672 slice dataset by intentionally introducing varying percentages of dropout noise. This dataset, with a median size of 4,015 spots, is well-suited for benchmark validation, as both DenoiseST and GraphST exhibit high Adjusted Rand Index (ARI) scores. Considering that dropouts are more prone to occur in genes with low expression levels, we introduced dropout noise by converting varying percentages of non-zero values (UMI count ≤ 5) to zero. For each specified percentage, we generate 10 random replicates, calculating the median and mean ARI scores. We observe that DenoiseST exhibits high stability even with small proportions of dropouts (10% and 20%). It continues to produce high ARI scores with an increase to 30% dropouts (**Figure 4B** and **Supplementary Table S3-S8**). Meanwhile, we replicated the analysis of artificial dropout noise for five alternative methods (GraphST, STAGATE, DeepST, SpaGCN, and BayesSpace), affirming that DenoiseST exhibits superior performance among them. Subsequently, we executed 10 random replicates for each percentage and computed the average ARI scores. This approach offers a more precise representation of variations in clustering accuracy.

### DenoiseST accurately identifies functionally variable genes with different spatial resolution

In SpaGCN, the concept of spatially variable genes (SVGs) has gained popularity. SpaGCN clusters the spatial domain and identifies SVGs for each spatial domain. To accurately identify SVGs, clustering accuracy is crucial, but SpaGCN cannot always cluster genes with perfect agreement with manual annotation labels. Moreover, the marker gene in spatial transcriptome data does not necessarily strictly follow the division of the manual annotation cluster distribution. The marker gene might be expressed in multiple clusters or might not be expressed in only one cluster. Therefore, we propose functionally variable genes (FVGs). To further verify the significance of the identified functionally variable genes, we specifically analyzed them in 151673 slice. First, we perform enrichment analysis on the Differential Expression Genes (DEGs) found by fitting different distributions for each cluster in the slice. We found that each cluster is enriched in pathways with certain differences (as shown in **Supplementary Figure S12 A**). Finally, the DEGs of each cluster are identified and counted (some outlier clusters may not be able to find the DEGs). If a DEG appears in at least 3(half of cluster number -1) clusters, we consider it a significant DEG.

We identify functionally variable genes through spatial continuity selecting among these significant DEGs. We performed GO enrichment analysis (**Figure 5B**) on the identified functionally variable genes and found enrichment in synapse organization and neuronal cells. This suggests that these genes may regulate the formation and maintenance of synapses^43^, signal transmission between pre-synaptic membranes and post-synaptic membranes^44^, synaptic plasticity^45^, etc., and may play an important role in the structure and function of neurons. DenoiseST(ARI=0.65) detected a total of 761 functionally variable genes (FVGs), while SpaGCN(ARI=0.38) identified 67 spatially variable genes (SVGs). We observed that SpaGCN’s gene selection process is overly stringent. We speculate that this phenomenon is primarily attributed to two reasons, namely, inaccurate clustering results obtained by SpaGCN. The distribution of gene expression in actual slices does not strictly align with the domain distribution of the ground truth, and a gene can be expressed in multiple domains. To validate our hypothesis, we examined the top 20 FVGs identified by DenoiseST (see **Supplementary Figure S13** and **Figure S14**). We conducted detailed analyses on SNCG, HOPX, YWHAG, and CA2 (**Figure 5A**). Our findings indicate that the SNCG gene exhibits high expression across layer 1, layer 2, layer 3, layer 4, and layer 5. The HOPX gene displays high expression in layer 1, layer 2, and layer 3. Similarly, the YWHAG gene shows high expression in layer 1, layer 2, layer 3, layer 4, layer 5, and layer 6. On the other hand, the CA2 gene exhibits higher expression in the WM layer.

**Figure 5.**
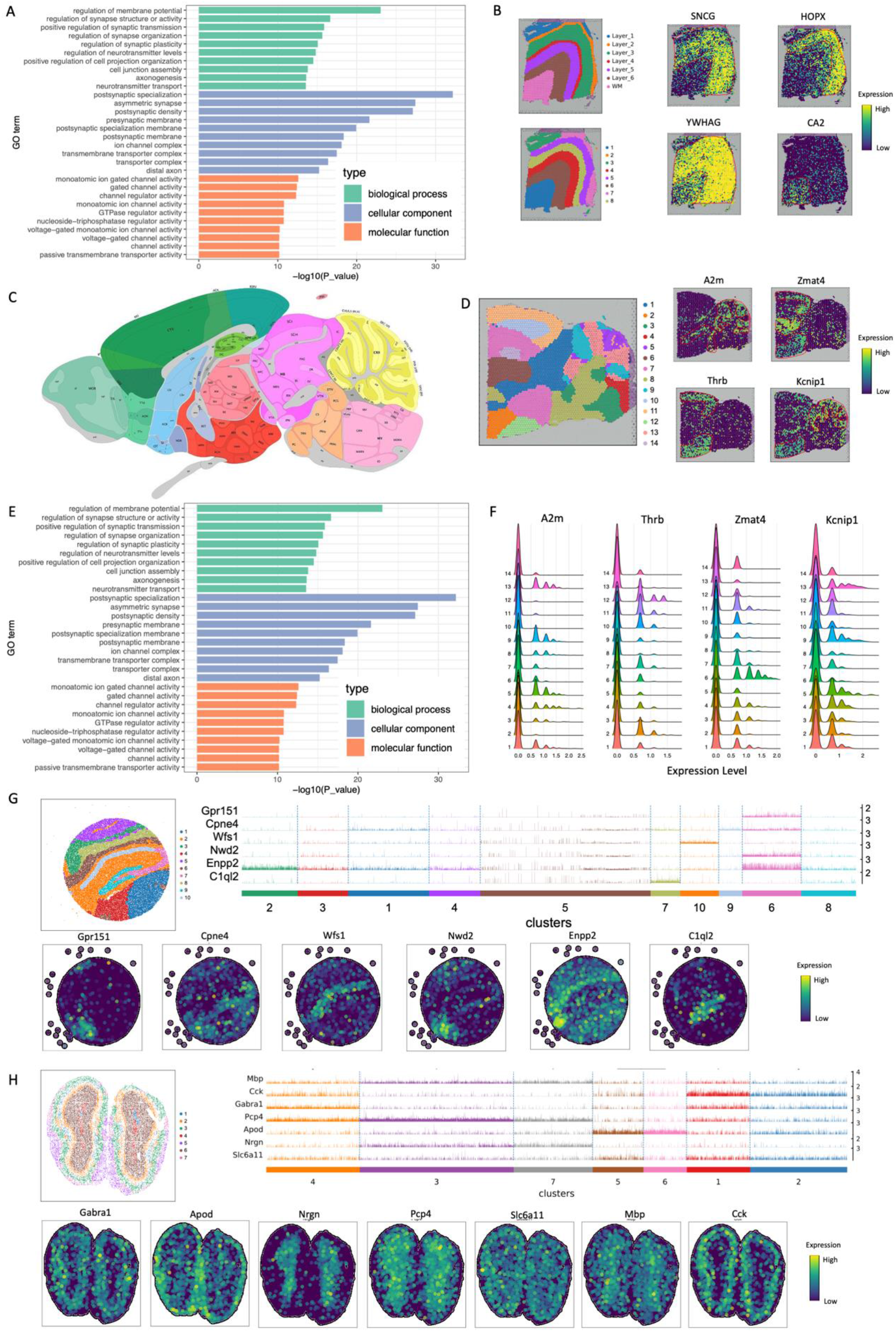
Functionally variable genes and high-resolution spatial transcriptome analysis. **(A)** Functionally variable genes (FVGs) detected in human dorsolateral prefrontal cortex 10x Visium dataset. **(B)** GO enrichment analysis in human dorsolateral prefrontal cortex 10x Visium dataset. **(C)** the annotated brain slice images from the Allen Mouse Brain Atlas. **(D)** GO enrichment analysis in mouse posterior brain slice. **(E)** GO enrichment analysis in mouse posterior brain slice. **(F)** Visualized the expression distribution of the A2m, Thrb, Zmat4, and Kcnip1 genes in each cluster on slices of the mouse posterior brain. **(G)** Visualization of spatial domains identified by DenoiseST and corresponding marker gene expression in mouse hippocampus tissue acquired with Slide-seqV2, utilizing these genes to generate heatmaps. **(H)** Visualization of spatial domains identified by DenoiseST and corresponding marker gene expression in mouse olfactory bulb Stereoseq data, utilizing these genes to generate heatmaps.

We analyzed the identified functionally variable genes (FVGs) in the mouse posterior brain slice and compared them with the annotated brain slice images from the Allen Mouse Brain Atlas (as shown in **Figure 5C** and **Supplementary Figure S13**). Each cluster in the slice undergoes enrichment analysis on the DEGs identified by fitting different distributions (as shown in **Supplementary Figure S12 B**). Every cluster demonstrates enrichment in pathways with distinct differences. Finally, we count the identified DEGs for each cluster. If a DEG appears in at least 6 clusters, we think it is a significant DEG.

Subsequently, we identify functionally variable genes among these significant DEGs. This analysis demonstrates that a gene can still be expressed in multiple domains. For example, the A2m gene and Kcnip1 gene can accurately identify regions composed of the pyramidal layer of the cortex and hippocampus that are anatomically similar in neuroanatomy. The high expression distribution of the Thrb gene and Zmat4 gene correlates with regions similar in structure to DenoiseST clustering results, such as domain11, domain2, domain12, and domain6. Furthermore, the GO enrichment analysis predominantly reveals enrichment in axonal synaptic tissue (**Figure 5E**). We believe that the functionally variable genes (FVGs) identified by our approach offer a more comprehensive consideration of both the gene expression matrix and spatial characteristics, resulting in higher research value. The manual annotation of ground truth in spatial transcriptomics slice data typically necessitates information such as marker gene expression, cell morphological characteristics, and cell function details. We think that the FVGs we have identified can assist in the precise identification of various cell types or cell subtypes, leveraging the gene distribution across different spatial domains. We also visualized the top 20 FVGs of the mouse anterior brain found by DenoiseST (**Figure 5D** and **Supplementary Figure S15**).

In **Supplementary Figure S12 C**, we statistically calculated and visualized the expression distribution of the SNCG, HOPX, YWHAG, and CA2 genes in each cluster on 151673 slices. In **Figure 5F**, we statistically calculated and visualized the expression distribution of the A2m, Thrb, Zmat4, and Kcnip1 genes in each cluster on slices of the mouse posterior brain. We observed distinct distributions of gene expression levels in each category. Notably, we found that the distribution states of these genes resemble the Poisson distribution, negative binomial distribution, and zero-inflated negative binomial distribution, respectively. This observation justifies our choice of fitting different distributions to the data when defining FVGs.

Finally, we evaluated the clustering performance of DenoiseST on high-resolution spatial transcriptomic data. Using Slide-seqV2 data (41,786 sub-cells) from the mouse hippocampus (**Figure 5G**), DenoiseST’s clustering results exhibited regional continuity. Heatmaps were generated for the expression levels of specific marker genes (Gpr151, Cpne4, Wfs1, Nwd2, Enpp2, C1qI2) found in this slice, and these were visualized on the slice. DenoiseST’s algorithm ensures that adjacent points belong to the same domain, providing stronger regional continuity. Furthermore, an analysis of the DenoiseST clustering results in the mouse hippocampus revealed that the cluster distribution aligns with specific marker genes. We also validated DenoiseST’s performance on Stereoseq chips of the mouse olfactory bulb (**Figure 5H**). Analyzing marker genes between DenoiseST domains, we identified specific laminally distributed genes (Gabra1, Apod, Nrgn, Pcp4, Slc16a11, Mbp, and Cck), consistent with previously reported assessments of genes in the mouse olfactory bulb dataset. DenoiseST’s clustering results reveal continuous regions with tissue textures similar to those observed in tissue slices (as shown in **Supplementary Figure S16 A** and **Figure S16 B**). Finally, we created heatmaps for these genes on all spots and visualized them on slices.

### DenoiseST-based biological significance analysis in breast cancer spatial transcriptomic data

To validate the capability of DenoiseST in discerning biological data, we employed DenoiseST to analyze a 10x Visium human breast cancer spatial transcriptomic dataset. This dataset was manually labeled by the pathologist, and these labels are considered the ground truth. Initially, we applied DenoiseST, STAGATE, and GraphST to cluster breast cancer dataset based on the equal cluster number with the ground truth (**Figure 6B**). Our findings indicate that DenoiseST acquires superior clustering accuracy. Considering that the pathologist’s annotation relies primarily on H&E staining and cell morphology, which overlooks transcriptomic variations, we employed DenoiseST to cluster this dataset in 30 clusters for a more detailed analysis.

**Figure 6.**
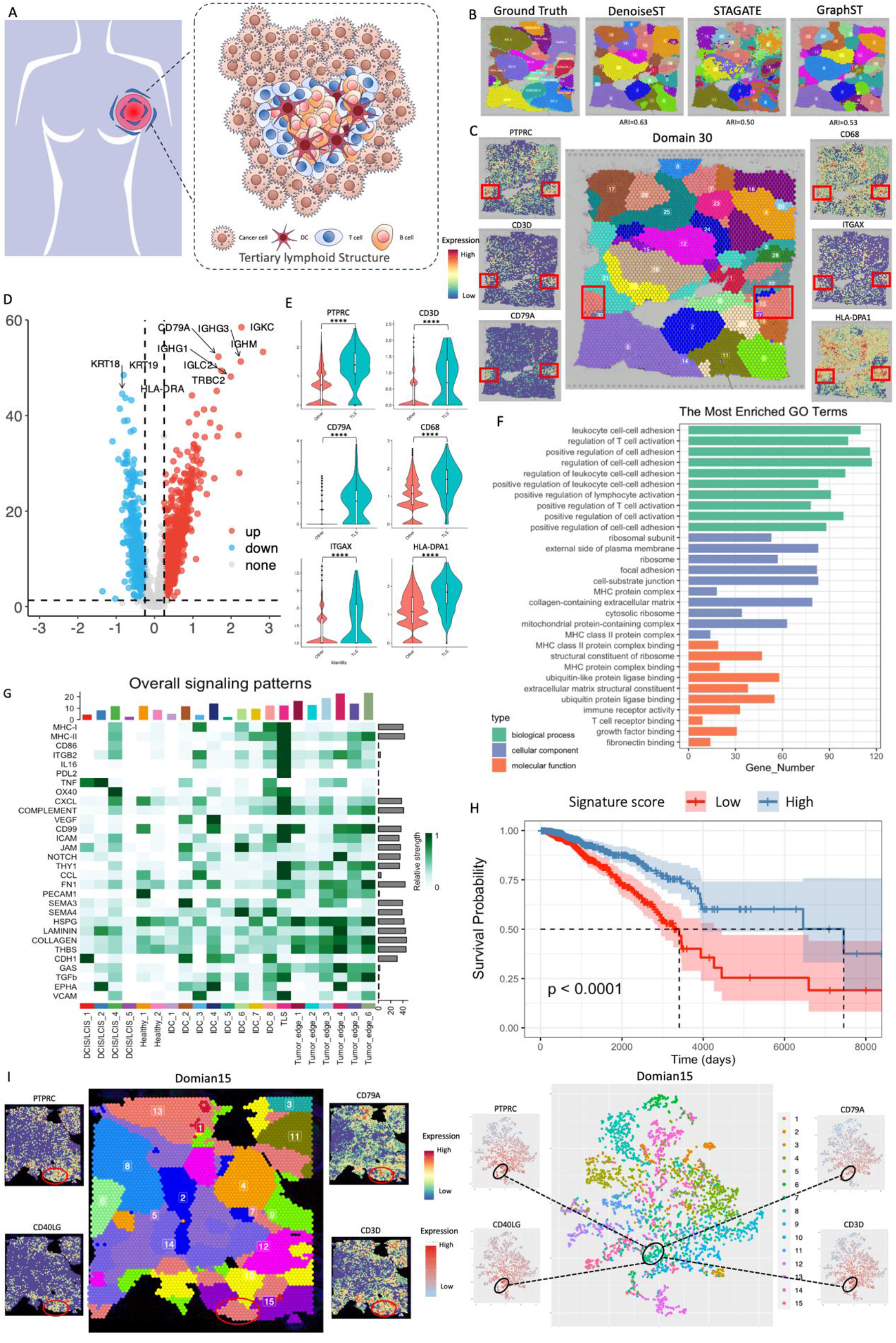
Analysis of biological significance in breast cancer spatial transcriptome data. **(A)** Schematic diagram of tertiary lymphoid structures (TLSs). **(B)** Comparison of clustering results in breast cancer slice 1 using DenoiseST, STAGATE, and GraphST. **(C)** Visualization of domain30 slice clustering results and TLSs-related genes for breast cancer slice 1. **(D)** Volcano plot of differentially expressed genes (DEGs) for cluster 13 of breast cancer slice 1. **(E)** Visualization of TLSs-related genes in cluster 13 and other regions. **(F)** GO analysis on cluster 13 of breast cancer slice 1. **(G)** Heatmap of enriched interaction pathways for breast cancer slice 1. **(H)** Survival analysis of TLS-signature score in TCGA-BRCA cohort. TLS-signature is constructed by gene markers from cluster 13 of breast cancer slice 1. **(I)** Visualization of domain15 slice clustering results and immune-related genes in breast cancer slice 2 and visualization of domain15 two-dimensional scatter plot clustering results for breast cancer slice 2.

Notably, in our analysis of the clustered results, we observed that cluster 13 exhibits a high expression of immune-related markers such as PTPRC, CD3D, CD79A, and ITGAX (**Figure 6C**). We also utilized the t-distributed stochastic neighbor embedding (t-SNE) method to visualize the gene expression matrix of spatial transcriptome data (**Supplementary Figure S17 A**). This expression profile suggests that the cluster is composed of various immune cells, including T cells, B cells, and dendritic cells^46,47^. Therefore, we hypothesize that cluster 13 represents tertiary lymphoid structures (TLSs). TLSs are specialized immune microenvironments formed in response to chronic inflammation or tumors^48^. Characterized by a dense composition of diverse immune cells such as T cells, B cells, and dendritic cells, TLSs play a vital role in orchestrating local immune responses (**Figure 6A**). To confirm our hypothesis, we conducted Differential Gene Expression (DEG) analysis on cluster 13, followed by Gene Ontology (GO) analysis (**Figure 6D, Figure 6E**, and **Figure 6F**). Differential Expression Genes (DEGs) such as IGKC, IGLC2, IGHG3, and CD79A were found to be highly expressed, which are typically associated with key immunological functions such as antibody production^49^, antigen presentation^49^, and lymphocyte activation^50^. These genes are indicative of active immune processes typically seen in TLSs. The GO analysis results further reinforced this, showing a significant association of cluster 13 with immune response pathways. These findings collectively support our hypothesis, suggesting that cluster 13 represents a tertiary lymphoid structure (TLS) within the tumor microenvironment.

Given that TLSs are composed of various immune cells and play a crucial role in the tumor environment, we conducted a cell-cell communication analysis to determine whether cluster 13 exhibits communication pathways characteristic of TLSs. As shown in **Figure 6G**, this analysis revealed the overall signaling patterns for each cluster. In cluster 13, we observed heightened activity in pathways such as MHC-I, MHC-II, CD86, ITGB2, IL16, and CXCL. This indicates a robust immune-related response, which is a hallmark of active TLSs. These findings lend strong support to our hypothesis that cluster 13 indeed represents a TLS.

TLSs are reported that associated with favorable prognosis in various cancers, including breast cancer. To further validate this in relation to cluster 13, we utilized the gene markers from this cluster to establish a signature score through Univariate Cox hazard analysis within the TCGA-BRCA dataset. We aimed to ascertain whether the signature score from cluster 13 correlates with improved patient prognosis. The results, as depicted in **Figure 6H**, showed that patients with higher signature scores from cluster 13 had a significantly better prognosis compared to those with lower scores. This finding aligns with the existing research that suggests a positive prognostic impact of TLSs in cancer. Taken together, the high expression of immune markers, active signaling pathways, and the prognostic significance of the signature score collectively affirm that cluster 13 is indeed representative of TLSs.

To further confirm DenoiseST’s robustness in elucidating biological datasets, we extended its application to another breast cancer dataset (**Figure 6I**). Remarkably, due to DenoiseST’s precise capability to discern biologically relevant clusters, we identified cluster 7, which exhibited high expression of gene markers such as PTPRC, CD79A, CD40LG, and CD3D. These markers are indicative of an area enriched with T cells and B cells. Validating our findings, DGE analysis was performed, with **Supplementary Figure S17 B** indicating that genes like TRBC2, TRAC, LTB, and IL7R were highly expressed in cluster 7. Further, GO analysis of the DEGs from cluster 7 highlighted immune-related pathways, particularly those associated with T cell and B cell functions, which suggests that cluster 7 is indeed enriched with these lymphocytes (**Supplementary Figure S18**). In conclusion, DenoiseST has demonstrated its ability to accurately identify biological structures within breast cancer datasets. Its effectiveness in revealing such structures emphasizes its potential to uncover critical biological insights from spatial transcriptomic data, which could significantly contribute to the advancement of targeted cancer treatments in the future.

## Discussion and Conclusion

Precise identification of spatial domains and the revelation of genes exhibiting functional variability play pivotal roles in comprehending tissue and biological functions. Here, we have developed the DenoiseST tool. The effectiveness of DenoiseST clustering is mainly attributed to four key aspects. First, DenoiseST uses a different feature set, helping to avoid the usage restrictions of a single feature set. Second, in the linear denoise model built by DenoiseST, a Pearson’s cell-cell similarity matrix is constructed, followed by transformations (PCA and Laplace transform) to reduce the negative impact of linear dropout noise. The resulting consensus matrix is then combined with the location information of the spatial transcriptome. The adjacency matrix is iterated, enhancing the affinity between adjacent cells to combat dropout noise. Third, in the nonlinear denoise model constructed by DenoiseST, the zero-inflated negative binomial and graph convolutional network model is utilized to interpolate the zero values of the original data and fuse the characteristics of adjacent cells, thereby countering the negative impact of nonlinear dropout noise in the data. Fourth, DenoiseST capitalizes on weight consensus clustering to merge multiple clustering results based on different feature sets. The incorporation of these four aspects fortifies DenoiseST’s robustness against dropout noise and mitigates the impact of false positives and false negatives in spot-spot connections. When clustering spots in the benchmark dataset, DenoiseST demonstrates superior clustering accuracy. Additionally, DenoiseST has designed a pipeline process that, based on the characteristics of the input spatial transcriptome data, adaptively models and conducts fully automated downstream analysis such as clustering. Choosing a modeling method suitable for the data helps reduce the risk of overfitting, enhances the model’s generalization ability, and improves modeling flexibility.

Spatial transcriptome identification of functionally variable genes poses a significant challenge due to the substantial amount of dropout noise and heterogeneity in the data. DEGman addresses this challenge by utilizing the Bhattacharya distance and testing multiple distributions. Specifically, DEGman employs the Bhattacharya distance and conducts tests across multiple distributions. Finally, histological information is employed for gene selection. In this study, our primary focus is on sequencing-based spatial transcriptomic (ST) data. We visualized the functionally variable genes (FVGs) identified in 151,673 slices of the DLPFC datasets and the mouse posterior brain slice. These FVGs, characterized by high spatial variability, underwent GO enrichment analysis, demonstrating enrichment in relevant tissue pathways. Notably, DenoiseST showed its capacity to process substantial amounts of high-resolution spatial transcriptome data.

In our analysis of human breast cancer data 1, we employed DenoiseST for clustering with a cluster number of 30 on breast cancer tissue slices. The clustering results exhibit clear boundaries and higher local aggregations. Notably, we found that cluster 13 is highly associated with tertiary lymphoid structures. Tertiary lymphoid structures play a crucial role in the immune system, contributing to the maintenance of immune function and suppression of tumor progression. We then validated cluster 13 through a series of in-depth analysis including DEG, GO, cell-cell communication analysis, and survival analysis. Additionally, in human breast cancer data 2, DenoiseST also identified the clusters with immune cell enrichment, which demonstrated the robustness of DenoiseST in its ability to analyze biological structures in spatial transcriptome data.

In this study, our primary focus is on low-resolution spatial transcriptomic data derived from next-generation sequencing. While DenoiseST can perform cluster analysis on high-resolution spatial transcriptome data, practical applications raise specific considerations. High-resolution spatial transcriptome data offers clearer image data. In algorithm design, it becomes crucial to not only incorporate the gene expression matrix and position information but also optimize the utilization of image information while avoiding noise integration. Additionally, the large size of data in high-resolution spatial transcriptome data necessitates careful consideration of both spatial and time complexity in algorithm design.

While DenoiseST demonstrates high performance in the analysis of 12 benchmark datasets, the application of this methodology necessitates careful consideration of several technical aspects. A key concern pertains to potential confounders introduced by batch effects. In the 12 benchmark datasets, each comprises three batches of slices. Batch effects across different datasets may obscure genuine biological signals^51^, with noise arising from disparate times, different processing personnel, and technological platforms, resulting in significant variations or batch effects. These effects can be either linear or nonlinear, making them challenging to distinguish from biological variability. Additionally, for the ST dataset with significant flaws, achieving the three-dimensional alignment of tissues remains a considerable challenge^17^. In such circumstances, addressing the integration issue of multiple batch data becomes imperative. In addition, the augmentation of DenoiseST’s capabilities can be realized through the integration of diverse single-cell omics datasets, including scRNA-seq, scATAC-seq^52^, and scMethyl-seq^53^. This integrative approach seeks to attain a holistic comprehension of cellular heterogeneity and epigenetic regulation. By accommodating both spatial transcriptome data and large volumes of single-cell RNA-seq data, we can gain a more comprehensive understanding of cellular diversity within tissues.

DenoiseST is designed to process spatial transcriptome data derived from next-generation sequencing and accommodate data from diverse experimental platforms. Validation of DenoiseST encompasses its application to 10x Visium, Slide-seqV2, and Stereo-seq datasets. Furthermore, DenoiseST is purposefully designed for computational efficiency, enabling effective handling of challenges posed by large datasets. Notably, testing on the 10x Visium dataset, comprising approximately 4,000 spots, demanded 20 minutes of wall-clock time on a server featuring an AMD EPYC 7502P 32-Core Processor CPU and NVIDIA Corporation GA100 GPU. As well as, the evaluation of a high-resolution dataset, comprising about 50,000 spots, required 30 minutes of wall-clock time on the same server configuration.

In conclusion, DenoiseST aims to mitigate the impact of dropout noise by employing both a linear denoise model and a nonlinear denoise model to address dropout noise and identify biological variations among cells. The versatility of DenoiseST is evident in its ability to handle datasets comprising tens of thousands of spots, showcasing promising outcomes across low-resolution and high-resolution datasets based on next-generation sequencing. To enhance computational efficiency, DenoiseST is endowed with a CPU+GPU heterogeneous parallel computing architecture, empowering users to analyze substantial datasets. In summary, DenoiseST has undergone testing as an effective method for clustering and identifying functionally variable genes, holding potential applicability across diverse realms of biological research by facilitating novel discoveries within spatial transcriptome data.

## Materials and Methods

### Methodology overview

The input dataset for DenoiseST consists of a gene expression counts matrix *M* and spatial positional information. In *M*, the columns correspond to spots, and the rows correspond to genes (referred to as features). Assuming that the set of spots is represented by *V* and the total number of spots is denoted as *n*, each element *m*_*ij*_ ∈ *M* signifies the expression level of the gene *i* in spot *v* _*j*_ ∈*V*. The workflow of DenoiseST is illustrated in **Supplementary Figure S1**. First, the selected gene subset (feature gene set) is separately input into both the linear denoise model and the nonlinear denoise model. To reduce feature extraction errors caused by individual feature sets, DenoiseST constructs multiple feature sets. The clustering results of these feature sets are obtained in the linear denoise model and the nonlinear denoise model respectively, and these clustering results are merged into a weight consensus matrix. Finally, spot types are determined through clustering analysis of the weight consensus matrix. The detailed description of the DenoiseST steps is provided below.

### Feature set selection and data preprocessing

We used the ‘seurat_v3’ function from the SCANPY^54^ and Seurat3^55^ packages to filter the raw gene expression data. The ‘seurat_v3’ function learns the mean-variance relationship from the data and calculates the mean and variance of each gene using unnormalized data. Then, a curve is fitted by computing a local polynomial fit of degree 2 to predict the variance of each gene as a function of its mean. This global fitting provides us with a regularized estimate of variance given the mean of a feature. Therefore, we can use it to normalize feature counts without removing changes higher than expected. Taking into account the anticipated variances, we executed the transformation

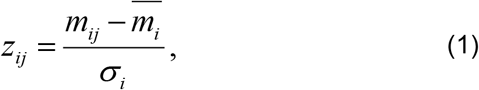

where *z*_*ij*_ represents the standardized value of the feature *i* in spot *j, m*_*ij*_ denotes the raw value of the feature *i* in spot *j*, 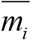signifies the mean raw value of features, and *σ* _*i*_ indicates the expected standard deviation of the feature *i* derived from the global mean-variance fit. The variance serves as a metric for measuring the dispersion of spot data after accounting for average expression. It is directly utilized for feature ranking and selection of the top gene features. Since it is challenging to determine the optimal number of top features for achieving the best clustering effect, a specific threshold (e.g., the top 3000 feature genes) is often set based on empirical knowledge. In this case, instead of using a single feature set, we utilized *fea* multiple feature sets that contain the top *T*_1_, *T*_2_, …, and top *T*_*fea*_ features.

Based on the evaluations of performance for various feature sets, *f*ea = 1,2,…,8 (see more details in **Supplementary Figure S10** and **Supplementary Figure S11**). After data analysis, each top *T*_*i*_ (*i* = 1,2,3) features’ gene set *G*_*i*_ typically does not exceed 25% of the total features in the dataset. For each selected set of feature genes 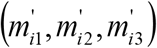, the log-transformed and normalized approach is carried out. Finally, these feature sets are inputted into the linear denoise model and the nonlinear denoise model, respectively.

### Linear denoise model

In the linear denoise model, we input the feature sets of different features sequentially.

#### Constructing spot similarity matrices

For a feature set *G*_1_ with selected *T*_1_ features, we keep *T*_1_ rows of features to obtain a *T*_1_ × *n* expression matrix *M*_1_. For any two spots *s*_*i*_, *s* _*j*_ ∈ *S*, the Pearson metrics *pea*(*s*_*i*_, *s*_*j*_) to construct distance matrices between *s*_*i*_ and *s* _*j*_ are calculated. *pea*(*s, s*) is constructed as follows:

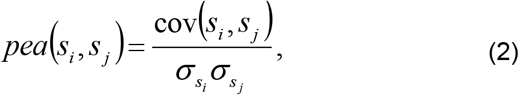

where cov(*s*_*i*_, *s*_*j*_) is the covariance of *s*_*i*_ and *s* _*j*_, 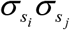 is the product of the standard deviations of *s*_*i*_ and *s* _*j*_.

We utilize the SC3^56^ package to convert the Pearson metrics through either principal component analysis (PCA) or by computing the eigenvectors of the related graph Laplacian. Subsequently, we arrange the columns of the resultant matrix in ascending order based on their corresponding eigenvalues. Next, we conduct k-means clustering on the first d eigenvectors of the transformed distance matrix (refer to **Supplementary Figure S1**). This is accomplished by employing the default k-means function in R and utilizing the Hartigan and Wong algorithm. Following that, we compute a consensus matrix *C* using the clustering-based similarity partitioning algorithm (CSPA)^57^. For each clustering result, we generate a binary similarity matrix based on the corresponding spot labels: if two spots belong to the same cluster, their similarity value is set to 1; otherwise, it is set to 0. To enhance computational efficiency, a consensus matrix is obtained by averaging the similarity matrices from each clustering. If the length of the *d* range of values (*D* in **Supplementary Figure S1**) exceeds 15, a random subset of 15 values is uniformly selected from the range and utilized.

#### Consensus matrix construction

Then, the consensus matrix *C*(*i, j*)_*n*×*n*_ is normalized, and the resulting normalized spot similarity matrix is as follows:

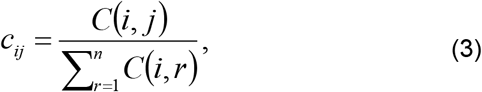

The normalized similarity matrix is defined by *C*_*nor*_ = (*c*)_*n*×*n*_, which is a *n*×*n* symmetric matrix and has 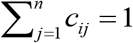.

DenoiseST transforms the spatial position information into an undirected adjacency matrix to improve the resemblance between a given spot and its neighboring spots. This process involves utilizing a predefined radius, denoted as *r*. Let *A* be the adjacency matrix, then, the value *A*_*ij*_ is set to 0.9 only if the Euclidean distance between spot *i* and spot *j* is smaller than *r* ; otherwise, *A*_*ij*_ is set to 0.1. The radius *r* is chosen empirically (the detailed list in **Supplementary Figure S10**).

#### Neighbor information enhancement and spectral clustering

Given a set of *fea* selected features, we can obtain the adjacency matrix *A*^(*l*)^ for the *l* th group of selected features by summing the normalized consensus matrix 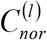. This process is performed for *l* = 1,2,3 to enhance the similarity between neighboring spots. The iteration follows the following formula:

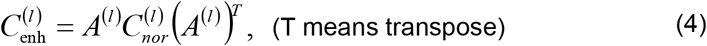

which can ensure that similarity information is exclusively propagated through the shared neighbors. For each final matrix 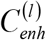, we utilize normalized spectral clustering^58^ by first calculating a normalized Laplacian 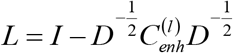 where *D* is defined as a diagonal matrix with the degrees (the total number of non-zero elements for each row of matrix 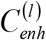) on the diagonal, then computes the first *k* eigenvectors of *L* to form a *n*× *k* matrix *U* and finally performs k-means clustering on the row-normalized matrix of *U*.

### Nonlinear denoise model

In the nonlinear denoise model, we input the feature sets of different features sequentially, resembling the linear denoising model.

#### Constructing graphs for spatial transcriptomics data

Spatial transcriptomics exhibits a robust association with spatial information, enabling efficient identification of similar cell states. To harness the full potential of this spatial information, we convert it into an undirected neighborhood graph *G* = (*V, E*), where each spot is connected to its predefined number of neighbors *k*. Within the graph *G, V* denotes a collection of spots, while E signifies a set of connected edges between spots. We represent the adjacency matrix of a graph *G* as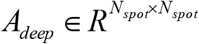, with *N*_*spot*_ indicating the total count of spots in the graph. If spot *j* ∈*V* is the neighbor of spot *i* ∈*V, a*_*ij*_ = 1, otherwise 0. Thus, for a given spot, its neighbors are determined by their proximity to other spots, which is computed using the Euclidean distance derived from the spatial location information. Finally, we select k spots from the top nearest neighbors as its neighbors.

#### Graph convolutional network

We design a graph convolutional network (GCN)^42^ based encoder to learn gene expression profiles and spatial location information. The encoder takes the neighborhood graph *G* of the neighborhood and the normalized gene expression matrix as input *X*_*m*_, and the decoder outputs the reconstructed gene expression matrix *H*_*s*_. To be specific, we employ a graph convolutional network (GCN) as an encoder to iteratively aggregate the representations of the neighbors and learn a latent representation *z*_*i*_ for a spot *i*. This process allows us to capture and incorporate the information from the neighboring spots to create a comprehensive and informative representation of each spot in the spatial transcriptomics data. Formally, the representations in the *lr* -th layer of the encoder can be expressed as follows:

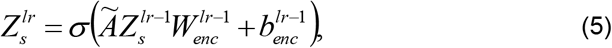

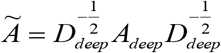 denotes the normalized adjacent matrix, wherein *D*_*deep*_ is a diagonal matrix having its diagonal elements. *W*_*enc*_ represents the training weight matrix, while *b*_*enc*_ denotes the bias vector. 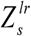 represents the output representation of the *lr* -th layer and 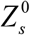 is designated as the original input gene expression matrix *X*_*m*_. *Z*_*s*_ is denoted as the final output of the encoder.

Subsequently, the latent representations *Z*_*s*_ are fed into a decoder to reverse them back into the raw gene expression space. In contrast to the encoder, the decoder adopts a symmetric architecture to reconstruct the gene expression. Specifically, the decoder is defined as follows:

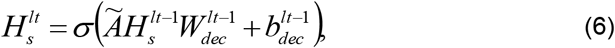

In this context, *H* represents the reconstructed gene expression profiles at the *lt* -th layer, while 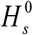 is initialized as the output representation *Z*_*s*_ of the encoder. *W*_*dec*_ and *b*_*dec*_ represent the trainable weight matrix and bias vector, respectively, which are shared by all nodes in the graph for the decoder. To maximize the utility of gene expression profiles, we train the model by minimizing the self-reconstruction loss of gene expressions through the following approach:

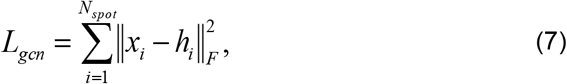

The output of the decoder results in denoted by *H*_*s*_, representing the reconstructed gene expression profiles. For spot *i, x*_*i*_ refers to the original normalized gene expression, while *h*_*i*_ corresponds to the reconstructed gene expression.

#### ZINB model-based graph convolutional network

To effectively simulate the distribution of spatial transcriptome data and acquire meaningful feature representations, we utilize a graph convolutional network based on the ZINB^30^ model. The ZINB distribution proves advantageous in capturing highly sparse and overdispersed gene expression data, allowing for more accurate and comprehensive modeling of the dataset.

The graph convolutional network based on the ZINB model, incorporates three separate fully connected layers that are connected to the final layer of the decoding layer. This architecture is employed to estimate the three essential parameters of the ZINB distribution: the dropout rate *π*, the dispersion degree *θ*, and the mean *μ*. To facilitate the calculations, we can express the parameters *π, θ*, and *μ* in matrix form. Let’s define their matrix representations as follows:

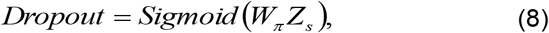

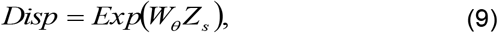

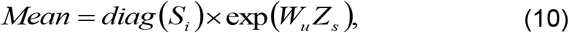

where *Z*_*s*_ = *f*_*dec*_ (*f*_*enc*_ (*X*_*m*_)) represents the output matrix obtained from the last layer of the decoding layer, while the size factor *S*_*i*_ is defined as the ratio of the total cell count to the median *S*. The ZINB model, which simulates the distribution of spatial transcriptome data, is parameterized by the ZINB distribution:

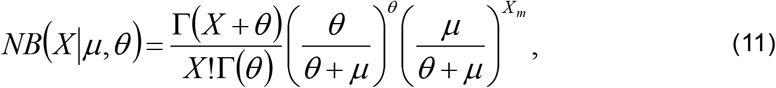

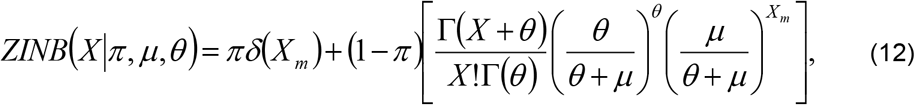

Ultimately, the loss function is defined as the sum of the negative logarithm of the ZINB distribution. The training objective of DenoiseST within this module is to minimize the loss function of the graph convolutional network based on the ZINB model:

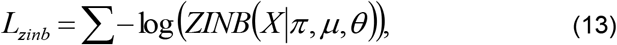

#### Overall loss function

The representation learning module for spatial transcriptome (ST) data is trained through the process of minimizing two distinct losses: the self-reconstruction loss and the ZINB loss. Briefly, the overall training loss of this model is defined as:

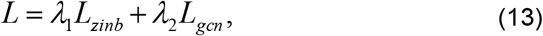

The weight factors *λ*_1_ and *λ*_2_ play a crucial role in balancing the impact of the reconstruction loss and the ZINB loss during training. Empirically, to achieve the desired trade-off between these two losses, we set *λ*_1_ to 1 and *λ*_2_ to 20.

#### Spatial domain clustering and refinement

After completing the model training process, the reconstructed spatial gene expression profiles *H* _*s*_ obtained from the decoder are utilized in conjunction with the nonspatial assignment algorithm mclust^59^ to cluster the spatial locations (spots) into distinct spatial domains. The mclust algorithm is commonly used for model-based clustering and is particularly effective when dealing with high-dimensional data, such as spatial transcriptomics data.

### Weight consensus clustering

The final result label of mclust clustering and spectral clustering for each *Lab*^(*l*)^ is saved as a *n*×*n* (0,1)-matrix, 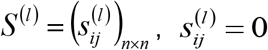 *or* 1, in which 1 and 0, respectively, represent that the corresponding two spots are and are not grouped.

Based on the *fea* matrices is constructed, where *S* ^(1)^, *S* ^(2)^, …, *S* ^(*f*)^, a weight consensus matrix *S* = (*S*_*ij*_)_*n*×*n*_ is constructed, where

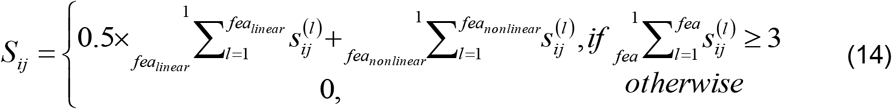

Proceeding with the same steps as before, we apply spectral clustering to the weight consensus matrix *S* in order to achieve the ultimate clustering result. Before applying to the cluster, we utilize the ‘Optimal_Clusters_KMeans’ function from the ClusterR^60^ package to automatically estimate the number of clusters. For low-resolution spatial transcriptomic data, we choose different models based on the dropout noise ratio. When the dropout noise ratio exceeds 90%, we apply both the linear denoise model and nonlinear denoise model to denoise the data, followed by weight consensus clustering. When the dropout noise ratio is less than 90%, to prevent overfitting of clustering results, we utilize the top 5000 feature genes in the nonlinear denoise model for data denoising, followed by clustering using ‘mclust’.

For high-resolution spatial transcriptomic data, to streamline the process and enhance computational efficiency, we employ the top 4000 feature genes in the nonlinear denoise model for data denoising, followed by clustering using Louvain or Leiden.

### Identifying functionally variable genes

The process of identifying functionally variable genes involves first recognizing differential expression genes and then, based on the characteristics of spatial transcriptome data, defining functionally variable genes.

#### Identifying differential expression genes in the spatial transcriptome

We employed the DEGman^61^ package to identify Differential Expression Genes (DEGs) in spatial transcriptome data with significant heterogeneity and data loss. Our approach involved optimizing the Bhattacharyya distance^62^ for three frequently used distributions (Negative Binomial^63^, Zero-Inflated Negative Binomial^64^, and Poisson^65^) through a combined strategy. Initially, we rapidly filter out genes without significant differences by employing the Bhattacharyya distance, a metric used to gauge the similarity between two probability distributions. This step helps us focus on genes that exhibit meaningful variations. Next, for the selected genes, we model their expression levels using three distributions: Negative Binomial (NB), Zero-Inflated Negative Binomial (ZINB), and Poisson. This process helps us determine the most suitable distribution for each gene. Subsequently, we calculate the Bhattacharyya distance between the best-fitting distributions of the two experimental conditions and perform a permutation test.

To obtain the confidence of scores of Bhattacharyya distance, we performed permutation tests to calculate *p* − *values*. The null hypothesis was that there was no difference in the expression distribution of each gene in the two spot groups. To test this, we performed *K* permutations where all cell columns were randomly shuffled. Then, the shuffled cells were sequentially divided into two groups of *m*_1_ and *m*_2_ spots. For each permutation, we calculated the Bhattacharya distance between the two spot groups for each gene. For a gene, given a distance score *B* between groups of original spots and K distance scores for K permutations (*B*_1_, *B*_2_, …, *B*_*K*_), the *p* − *value* for that gene is calculated as

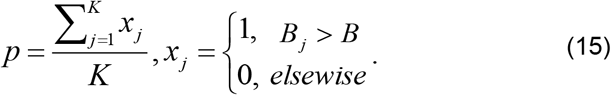

To further refine our selection and exclude genes whose expression distribution did not show significant differences under a predefined threshold, we employed the “p.adjust” function with the “FDR” parameter for *p* − *values* adjustment using False Discovery Rate (FDR) control^66^. The genes with adjusted *p* − *values* under 0.05 were then identified and chosen as the set of significant DEGs.

#### Identifying functionally variable genes in the spatial transcriptome

To identify more prominent marker genes, we define them as functionally variable genes. We then take the intersection of marker genes defined by each cluster. For example, if there are 7 clusters, we retain gene names that appear in more than 3 clusters. Subsequently, we mapped the gene expression onto each spatial spot and chose the top 100 spots for each gene with the highest expression levels. Then, we computed the distance between each spot, summing their count if the distance fell below a specified threshold. A higher numerical value suggests greater spatial continuity, indicating a higher likelihood of histological relevance, and categorizes the gene as a functional variable (Refer to **Supplementary Note S2** for specific details).

## Author Contributions

Leyi Wei and Xiucai Ye conceived and supervised this project. Yaxuan Cui designed the DenoiseST model, and Yaxuan Cui and Ruheng Wang built the model. Yaxuan Cui, and Yang Cui wrote the manuscript. Xin Zeng and Yang Cui led the biological data analysis. Yaxuan Cui, and Zheyong Zhu assembled the thesis pictures. Yaxuan Cui and Ruheng Wang revised the paper.

## Availability of Data and Materials

The source code package is freely available at https://github.com/cuiyaxuan/DenoiseST/tree/master. The datasets used in this study can be found at https://drive.google.com/drive/folders/1H-ymfCqlDR1wpMRX-bCewAjG5nOrIF51?usp=sharing. The details of data description are shown in **Supplementary Note S1**.

